# Targeting antibiotic resistant bacteria with phages reduces bacterial density in an insect host

**DOI:** 10.1101/493718

**Authors:** Lauri Mikonranta, Angus Buckling, Matti Jalasvuori, Ben Raymond

**Affiliations:** University of Exeter, Biosciences, Penryn Campus, Penryn, Cornwall, TR10 9FE, UK.; University of York, Department of Biology, Wentworth Way, York, YO10 5DD, UK.; University of Jyväskylä, Department of Biological and Environmental Science, Nanoscience Center, PL 35, 40014, Jyväskylä, Finland

**Keywords:** antibiotic resistance, bacteriophage, Enterobacter cloacae, gut infection, insect model, phage therapy

## Abstract

Phage therapy is attracting growing interest among clinicians as antibiotic resistance continues becoming harder to control. However, clinical trials and animal model studies on bacteriophage treatment are still scarce and results on the efficacy vary. Recent research suggests that using traditional antimicrobials in concert with phage could have desirable synergistic effects that hinder the evolution of resistance. Here, we present a novel insect gut model to study phage-antibiotic interaction in a system where antibiotic resistance initially exists in very low frequency and phage specifically targets the resistance bearing cells. We demonstrate that while phage therapy could not reduce the frequency of target bacteria in the population during positive selection by antibiotics, it alleviated the antibiotic induced blooming by lowering the overall load of resistant cells. The highly structured gut environment had pharmacokinetic effects on both phage and antibiotic dynamics compared to *in vitro*: antibiotics did not reduce the overall amount of bacteria, demonstrating a simple turnover of gut flora from non-resistant to resistant population with little cost. The results imply moderate potential for using phage as an aid to target antibiotic resistant gut infections, and question the usefulness of *in vitro* inferences.

## BACKGROUND

Problems arising from antibiotic resistant infections are set to increase worldwide. As a consequence, bacteriophages (bacteria-specific viruses) have started to attract serious consideration as antimicrobial agents after this approach was largely forgotten for decades in the Western world [1, 2]. However, their efficacy as therapeutic agents remains controversial [3]. Some of the *in vivo* work promises therapeutic potential in insect [4], mouse [5], and human infections [6], and *in vitro* experiments show overwhelming evidence of phages controlling the population size of their host bacteria [7]. Despite this, the few existing modern and properly controlled medical trials report varying success [3, 6, 8]. Key limitations of phage therapy include high specificity, ease at which bacteria can evolve resistance, and localised activity in the body.

It has been argued that phages may be of particular therapeutic value when combined with antibiotics, by constraining the emergence and spread of antibiotic resistance [9]. A good example of recent success in combination treatment comes from a difficult chronic *Pseudomonas aeruginosa* infection in human aortic graft [10]. Theoretically, phages may limit antibiotic resistance during treatment because of reduction in population size and synergistic costs of resistance [9]. Moreover, with cases where antibiotic resistance is carried on plasmids, phages can be used to directly target the plasmid carrying cells [11]. However, this latter approach has only been investigated *in vitro* [11, 12], which ignores a plethora of selection pressures towards both the host bacteria and the phage, such as host immune system, spatial structures within the tissues, nutrient availability and presence of native microbial flora [13].

Here, we studied the effects of antibiotic and phage treatment to bacterial load and frequency of resistant cells in a gnotobiotic insect gut model system, where phages target the bacteria harbouring antibiotic resistance plasmids. We compare these findings to *in vitro* context to assess the usefulness of inferring *in vivo* dynamics from *in vitro* studies. A full factorial setup of antibiotics and phage were orally administered to cabbage looper (*Trichoplusia ni*) larvae harbouring *Enterobacter cloacae* gut bacteria and a low initial frequency of antibiotic resistance plasmid. We show that targeting the tetracycline resistant cells with phage could not prevent the increase in resistance frequency in the presence of antibiotics. However, phage reduced the bacterial population size when plasmid was driven to high frequency with antibiotic selection.

## MATERIALS AND METHODS

*Enterobacter cloacae* wild type strain, that forms persistent gut association with Lepidopteran larvae (e.g. *P. xylostella, Mamestra brassicae and Trichoplusia ni)* after oral inoculation, was isolated from *Plutella xylostella* in the insectary of Department of Zoology, University of Oxford. Two spontaneous antibiotic resistant mutants, ANC C2 (rif^R^) and 11.1B (strep^R^ + nal^R^) were used in this experiment. The IncP-type plasmid RP4, coding for tetracycline resistance, is usually conjugative [14], and the lytic, plasmid dependent tectivirus PRD1 specifically infects RP4-bearing cells by recognising the bacterial sex-apparatus [15]. However, in this system the plasmid is essentially non-conjugative and functions as a carrier of tetracycline resistance and phage receptor genes. The mechanistic basis for the poor RP4 conjugation in these *E. cloacae* strains is unclear. Given that the phage is still infective, it could be that ANC C2 rifampicin-resistant mutant was an impotent recipient for the plasmid rather than that 11.1B(RP4) lacked the ability to express pili. Although rifampicin resistance mutations are most often polymerase related, known mechanisms in *Enterobacteriaceae* include elongated or abundant outer membrane LPS-chains [16] that could theoretically interfere with conjugation (but see: [17]).

The *in vitro* experiment used 200 μl cultures on a 96-well plate with LB and 0.1% starting frequency of plasmid. 24 replicate populations were subjected to full factorial antibiotic (AB) and phage (P) treatment: AB− P−, AB− P+, AB+ P−, and AB+ P+, respectively. The antibiotic treatments were set to 12 μg/mL tetracycline and phage starting density to 5 × 10^7^ phage particles. The growth in optical density (600nm) was recorded at 24 h with a spectrophotometer in 30°C. Plasmid frequency and transconjugants were sampled from a subset (N=44) of populations at 24 h by selective plating.

The *T.ni* were maintained as follows: The adults, fed with 100mM sucrose solution, were let to mate and lay eggs on paper strips in flight chambers. The eggs were placed in autoclaved, single-filter vented, 305 × 203 × 100mm Genesis™ containers for surgical instruments to maintain sterile conditions while allowing gas exchange. Hatched larvae were let to feed on Hoffman diet (288 g wheatgerm, 132 g caesin, 117 g sucrose, 70 g agar, 57 g dried brewers yeast, 37.5 g Wesson’s salts, 6 g sorbic acid, 3.75 g cholesterol, 3.75 g methyl-4-hydroxybenzoate and 2 g Vanderzant vitamin mixture in 2750 ml water) [18]. The liquid medium was poured into a sterile steel mould and solidified to diet cubes with approximately 25mm edges. The cubes were placed on Petri dish covers inside the containers. All rearing and *in vivo* bacterial work was carried out in 25 °C. For population maintenance, 50-60 pupae were removed from their cocoons per flight chamber and let to emerge and mate to start a new generation of hosts.

For inoculating the larval gut with *E. cloacae*, clones ANC C2 and 11.1B bearing RP4 were grown over night in LB (30 °C, 180 rpm shaking), the latter diluted 1:10 with 0.85 % NaCl, and then mixed together in a 50:1 ratio. The frequency of the plasmid bearing cells was counted by dilution plating as 0.072% and bacterial density approximately 1.2 × 10^8^ CFU. Diet cubes were dipped in this solution, dried for 5 min, and 3^rd^ instar larvae were allowed to feed on this bacterial diet for 30 h. The larvae were then subjected to the AB− P−, AB− P+, AB+ P−, and AB+ P+ treatments. Tetracycline (200 μg/mL final concentration) was added directly to the autoclaved diet mix and 30 μl of 10^10^ pfu/mL filtered phage lysate was pipetted onto each face of the phage treatment diet cubes. The experiment was carried out in two independent batches (final N=156 larvae). After 96 h, the larvae were surface sterilised with 70% alcohol and homogenised in 500 μl 0.85% NaCl with Qiagen Tissuelyser II™. The homogenate was serially diluted and plated on 12 μg/mL tetracycline, 12 μg/mL tetracycline + 100 μg/mL rifampicin, and LB plates to calculate the frequencies of plasmid carriers, transconjugants, and total density of bacteria. 24 bacterial clones from the phage treatment serial dilutions on tetracycline were tested for resistance against the ancestral phage with a phage plaque overlay assay.

Phage survival in the gut was confirmed by overlaying homogenised larval faecal samples on ancestral 11.1B(RP4 lawn on soft agar and observing plaque formation (N=24).

Treatment effects on densities and frequencies in both experiments were analysed with SPSS statistics 21.0 with antibiotics, phage, and antibiotics*phage interaction in ANOVA-GLM model. Post-hoc multiple comparisons were Bonferroni corrected.

## RESULTS

*In vitro*, antibiotics and phage had synergistic effects on the total density of bacteria at 24h (AB: F_1, 96_=3528.0, p<0.001, P: F_1,96_=73.7, p<0.001, AB*P: F_1, 96_=97.0, p<0.001). Multiple comparisons showed that there was no phage effect in absence of antibiotics but other pairwise differences were highly significant (p<0.001, Fig. 1. a). Thus, single treatment allowed compensatory growth of the non-target strain, but this effect was reduced for the plasmid carriers in the presence of antibiotics. Contrary to our expectations and previous studies with the same plasmid and phage in *E. coli* [12], the plasmid went to near fixation in the presence of antibiotics (F_1, 44_=8864.4, p<0.001) regardless of the phage (P and AB*P: p>0.05). This is particularly surprising, as unlike in *E. coli*, plasmid RP4 had very low rates of conjugation in *E. cloacae*: transconjugants were present in only one of the 44 sampled *in vitro* populations (AB+P− treatment at a very low frequency, while the phage remained infective. The cost of plasmid carriage in the absence of antibiotics manifested itself as complete plasmid extinction (at least below the detection threshold of <67 cfu/mL) in all sampled populations (Fig 1. b). This was the case even though RP4 has been reported to be of low cost and rapidly evolve towards no cost through chromosomal compensatory mutations [19].

**Figure 1.**
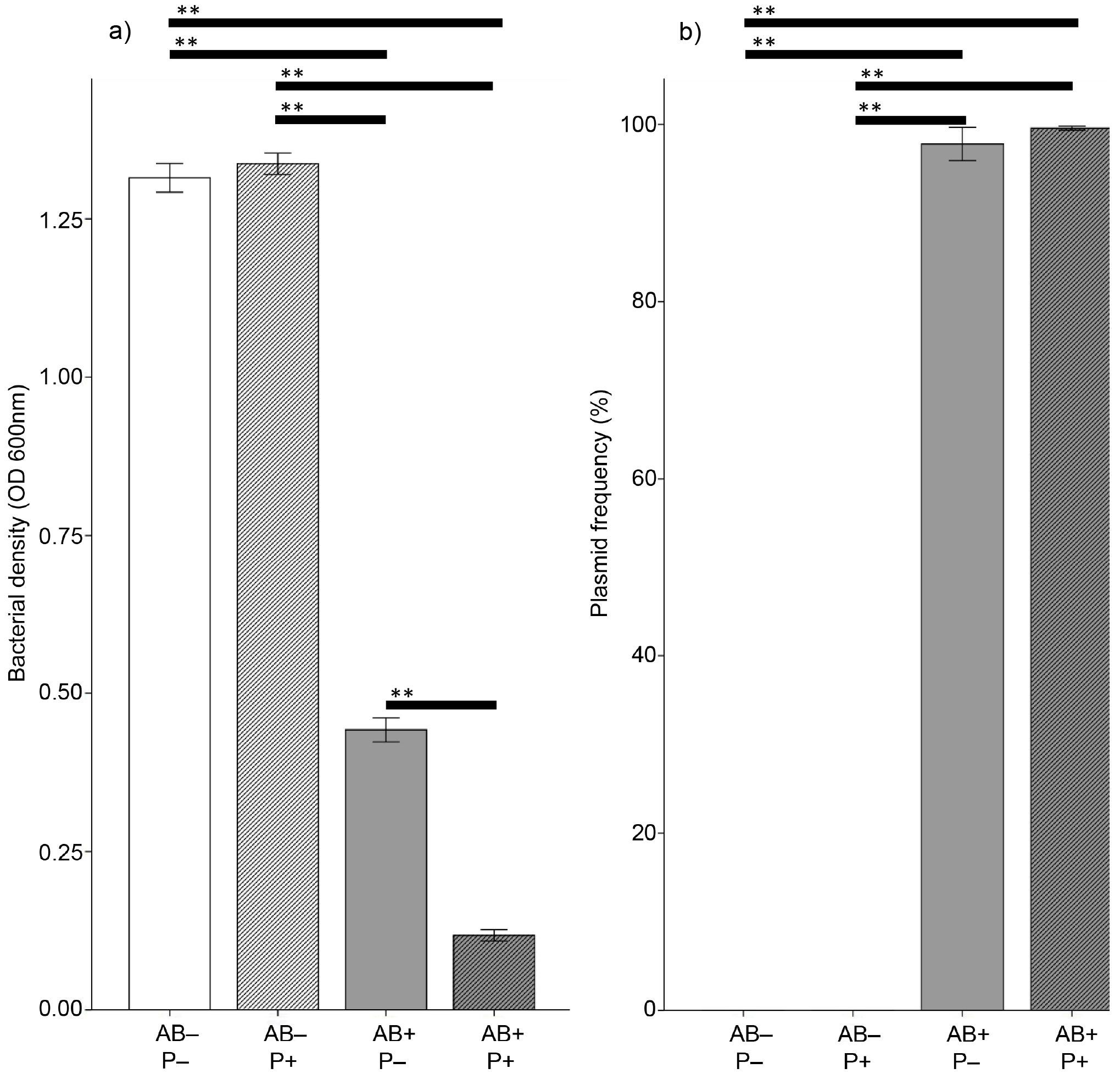
Dynamics in vitro. The density of bacteria (a), and the frequency of plasmid bearing cells (b) in liquid culture after a 24 h factorial antibiotic (AB) and phage (P) treatments. Error bars are +/− 1 S.E., (**) denotes p<0.001 difference.

*In vivo* patterns of plasmid dynamics were mainly similar to that observed *in vitro,* but the effects smaller. The non-targeted strain could fully compensate for the reduction in density of the competing strain (AB: F_1, 156_=0.1, p=0.77), and phage had no main effect on total bacterial load (P: F_1, 156_=3.0, p=0.085). Phage however lowered bacterial density in the presence of antibiotics (interaction: F_1, 156_=6.7, p=0.011), demonstrating that the treatment synergism was maintained *in vivo* (post hoc AB+ P− and AB+ P+, p=0.020, Fig. 2. a). Plasmid increased in frequency in all treatments but tetracycline selection drove the majority of the population to be plasmid carriers with no effects of phage addition (effects on plasmid frequency, AB: F_1, 156_=845.0, p<0.001; P: F_1, 156_=0.6, p=0.43; AB*P: F_1, 156_=0.4, p=0.55 (Fig. 2. b). Thus, phage therapy could not restrict the spread of antibiotic resistance in the *T. ni* gut.

**Figure 2.**
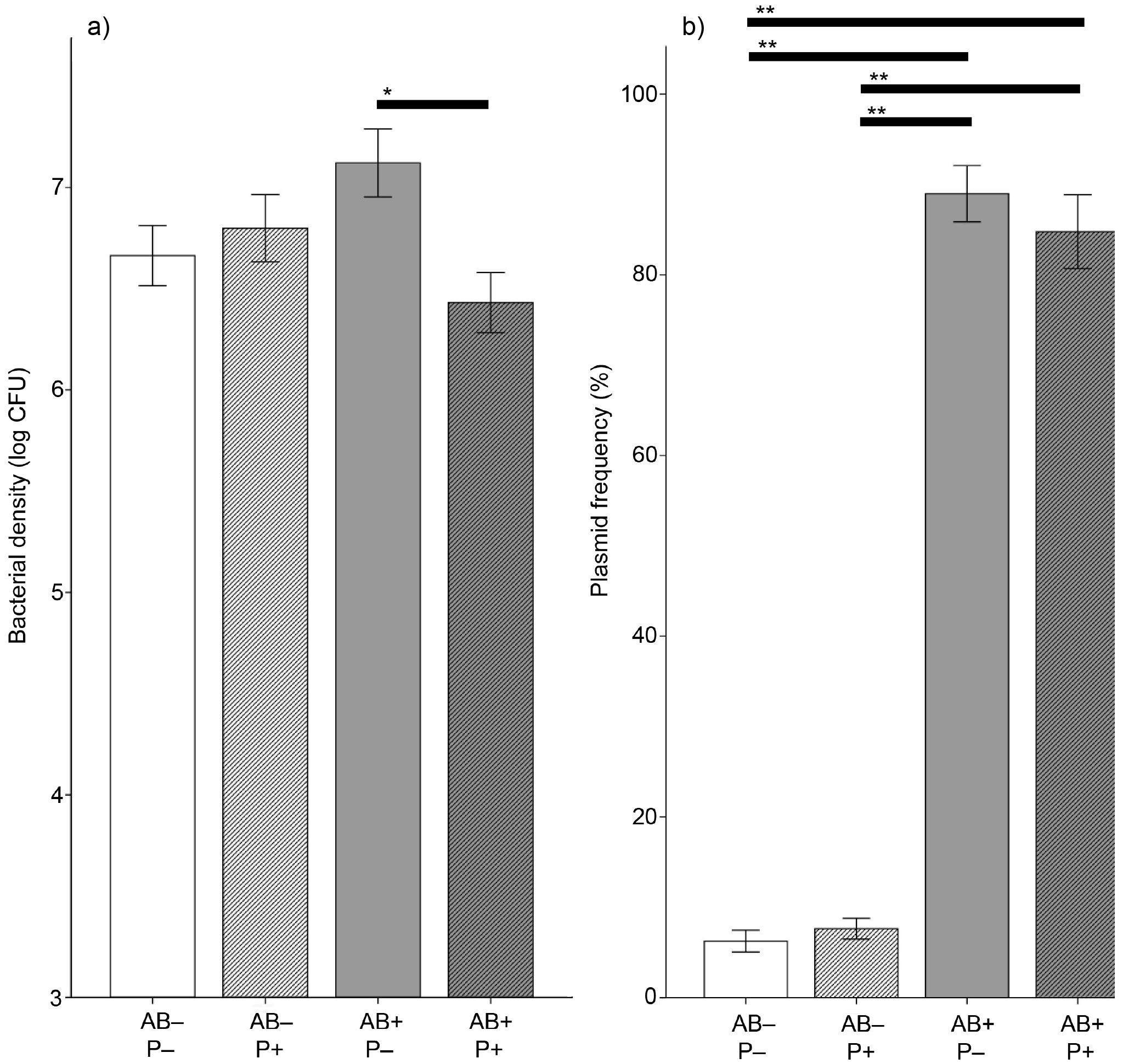
Dynamics in vivo. The overall density of bacteria (a) and the frequency of plasmid bearing cells (b) in the larval guts after a 96 h factorial antibiotic (AB) and phage (P) treatments. Error bars are +/− 1 S.E. (*) and (**) denote p<0.05 and p<0.001 differences, respectively.

## DISCUSSION

Here, we investigated the effect of plasmid-targeting phage on the spread of antibiotic resistance plasmid *in vitro* and in an insect gut model. Generally, the complex environment in the gut made treatment effects less pronounced compared to the liquid culture: apparently the spatial structures protected the plasmid from extinction in the absence of antibiotics, and allowed more non-resistant bacteria to coexist in the presence of tetracycline. This happened even though the antibiotic concentration in the diet was over 16 times higher than in the liquid culture and the experiment length 4 times longer, allowing more time for potential fixation and/or extinction to happen. Phage driven reduction in the bacterial population size in the presence of tetracycline was also more moderate *in vivo* but it is also notable that no detectable phage resistance emerged in the sampled populations. Surprisingly, antibiotics did not reduce the overall bacterial density in the gut, but rather replaced non-resistant cells with resistant ones while population size remained the same. In other words, antibiotics increased the relative fitness of the resistant strain without reducing the absolute fitness [20]. This was in contrast with liquid culture where the plasmid went even closer to fixation in the presence of antibiotics but resulted in lower total bacterial density, which suggests a greater growth cost associated with antibiotic resistance *in vitro*. We cannot attribute the inversion of fitness effects between *in vivo* and *in vitro* conditions (increase in plasmid frequency *in vivo*, and extinction *in vitro* in absence of antibiotics directly to the plasmid, because it could also be due to the differences in the mutant strains. The result still highlights the limitations of making inferences from test tube dynamics.

In the absence of antibiotics neither *in vitro* nor *in vivo* regimes showed a phage effect on the bacterial population size, which was not surprising because of very low initial frequency of phage-susceptible (i.e. plasmid bearing) cells. Most importantly, when antibiotic selection increased the frequency of phage-susceptible cells by spreading antibiotic resistance, phage decreased the total density of bacteria. This effect was bigger *in vitro* but also clearly visible in the insect gut. The gut lumen is a favourable environment for bacterial biofilm formation [21], which can offer non-resistant cells spatial protection from antibiotics [22] and/or from phage [23]. On the other hand, it could be that the observed phage efficacy to reduce bacterial load *in vivo* is partly due to their known ability to eradicate biofilms [24, 25], which are essential for bacterial persistence in the insect gut [21].

One potential epidemiological implication of viable phages in the faeces is that the viral therapeutic agent could restrict ongoing pathogen transmission or be transmitted with the target pathogen [26]. The model system presented in this study allows the manipulation of structure and connectivity of the population of the treated organism and would be an interesting future endeavour towards epidemiological effects of phage therapy. Our findings have implications for therapeutic use of phage, especially considering synergistic evolutionary effects with antibiotics [9]. However, antibiotics overwhelmed the selection imposed by the phages, meaning that the relative effect remained rather small. This could be improved with using more efficient phages or phage cocktails in a way that can target wider niche space *in vivo* by, for example, biofilm specificity [25]. It has also been shown that the effect of phage on plasmid-maintenance can depend on the type of antibiotics that are used [27]. To conclude, within-organism pharmacokinetics and the spatial, temporal, and resource-related dynamics are very different compared to a test tube, having major effects on cost of resistance, magnitude of selection, and treatment efficacy.

## ACKNOWLEDGEMENTS

The authors thank Alex Robinson for assistance with *T.ni* rearing.

## AUTHOR CONTRIBUTIONS

All authors conceived the study and designed the experiments. LM did the experimental work and analysed the data. All authors contributed to writing and approved the final version of the manuscript.

## ETHICAL STATEMENT

N/A

## DATA ACCESSIBILITY

The data will be made accessible in Dryad upon acceptance.

